# Data-driven analysis of facial thermal responses to an emotional movie reveals consistent stimulus-locked physiological changes

**DOI:** 10.1101/2020.09.02.276592

**Authors:** Saurabh Sonkusare, Michael Breakspear, Tianji Pang, Vinh Thai Nguyen, Sascha Frydman, Christine Cong Guo, Matthew J. Aburn

## Abstract

Facial infra-red imaging (IRI) is a contact-free technique complimenting the traditional psychophysiological measures to characterize physiological profile. However, its full potential in affective research is arguably unmet due to the analytical challenges it poses. Here we acquired facial IRI data, facial expressions and traditional physiological recordings (heart rate and skin conductance) from healthy human subjects whilst they viewed a 20-minute-long unedited emotional movie. We present a novel application of motion correction and the results of spatial independent component analysis of the thermal data. Three distinct spatial components are recovered associated with the nose, the cheeks and a respiratory component. We first benchmark this methodology against a traditional region-of-interest based technique. We then show significant correlation of all the physiological responses across subjects, including the thermal signals, suggesting common dynamic shifts in emotional state induced by the movie. Finally, we show that thermal responses were significantly anti-correlated with the positive emotional content of the movie thus an index of emotionally-driven physiological response. In sum, this study introduces an innovative approach to analyse facial IRI data and highlights the potential of thermal imaging to robustly capture emotion-related changes in ecological contexts.

## Introduction

Measures of physiological correlates of emotion predominantly use galvanic skin response (GSR) and heart rate (HR) metrics (Cacioppo & Tassinary, 1990; Kreibig, 2010; Rainville, Bechara, Naqvi, & Damasio, 2006). However, it has been argued that measures targeting a variety of somatic effects may be best able to capture emotion-related brain-body states (Kreibig, 2010). An innovative technique which compliments the traditional measures is facial infra-red imaging (IRI). IRI quantifies the temperature fluctuations of the face resulting from blood flow changes (Ioannou, Gallese, & Merla, 2014). As a contact-free technique, it allows ecologically valid experimental conditions which recent studies have exploited (Srivastava et al., 2020). Although it has not yet been widely used, thermal imaging has shown promising results in affective research (Ebisch et al., 2012; Engert et al., 2014; Ioannou et al., 2013).

The current use of thermal imaging analysis is based on ad hoc choices of facial location, usually chosen *a priori* and used with a region of interest (ROI) analysis i.e. extracting the thermal signals from specific facial regions like the nose-tip, the cheeks and the forehead (Ebisch et al., 2012; Engert et al., 2014; Ioannou et al., 2013; Kuraoka & Nakamura, 2011; Pavlidis, Levine, & Baukol, 2001; Pinti, Cardone, & Merla, 2015). ROI-based thermal signal extraction can be strongly influenced by motion artefacts (Strakowska, Strakowski, Wiecek, & Strzelecki, 2012) with only a few studies implementing motion tracking or motion correction (Manini et al., 2013; Sonkusare, Ahmedt-Aristizabal, et al., 2019). Furthermore, ROI based approaches have their own unique challenges and limitations. First, they only exploit a small portion of the information encoded in the whole face. Second, manual steps are time consuming and only feasible for the analysis of small data sets with low sampling rate. Third, different choices for the size of ROIs, their shapes and placement can further introduce biases (Ioannou et al., 2014).

Although thermal signal extraction based on ROIs has provided many insights into thermal effects in affective research, these methods can be complemented by data-driven approaches which uncover underlying data features with less *a priori* assumptions. Independent component analysis (ICA), specifically spatial ICA (sICA), is a subset of such methods for blind signal separation employed under assumptions of statistical independence of the source signals (McKeown, Hansen, & Sejnowsk, 2003). Due to the effectiveness in capturing the essential structure of diverse kinds of data, ICA has been widely used in many applications, such as functional magnetic resonance imaging (fMRI) analysis (Beckmann, DeLuca, Devlin, & Smith, 2005; Calhoun, Adali, Hansen, Larsen, & Pekar, 2003) which measures the haemodynamic changes as a proxy for neuronal activity. Similar to the brain, facial skin has an extensive distribution of blood vessels. Furthermore, temperature changes associated with the face can have various contributions, including stimulus induced chances but also respiratory confound, perspiration, and noise. The diverse nature of these signals suggests that blind signal separation techniques could be advantageous for isolating the different sources of signal (McKeown et al., 2003) and distinct spatial components.

With these considerations in mind, we sought to employ a sICA approach to characterize the dynamic facial thermal responses during naturalistic emotional experience. For this we acquired facial IRI data alongside other traditional physiological measures while subjects watched an emotionally salient movie. Movies are relatively unconstrained, motivating the need for data-driven signal extraction approaches. To analyse thermal signals in a maximally naturalistic setting where participants were free to move, we combined state-of-the-art motion correction method based on optical flow with sICA to extract dynamic temperature changes that manifest in different facial regions. Since ROI-based facial IRI studies have recognized the nose-tip as the most sensitive region for temperature fluctuations, we hypothesised that a nose-tip spatial component would be reliably captured with the sICA method as well.

To induce affective states in subjects, we employed a long continuous and an unedited emotional movie (∼20 minutes). Whereas static stimuli such as photos have often been used with thermal imaging in the past (Salazar-López et al., 2015) they have limitations for evoking the physiological responses (Gross & Levenson, 1995) that are integral to emotional experiences (James, 1922). Naturalistic stimuli have emerged as an alternative to strictly controlled paradigms and stimuli, such as pictures and sounds, as they offer better ecological validity (Hasson, Nir, Levy, Fuhrmann, & Malach, 2004; Sonkusare, Breakspear, & Guo, 2019). In neuroimaging studies, inter-subject correlation (ISC) analyses of movie viewing data have demonstrated common covarying patterns of brain activity (Hasson et al., 2004; Nguyen et al., 2016; Pajula, Kauppi, & Tohka, 2012). However, is the shared quality of affective experience between people detectable at the physiological level? Here we investigate whether such shared responses are present in various physiological signals (heart rate metrics, skin conductance and thermal responses). Furthermore, reports also suggest the differentiating role of facial temperature in emotional valence [(Matsukawa et al., 2017; Zenju, Nozawa, Tanaka, & Ide, 2004) but also see (Kosonogov et al., 2017; Salazar-López et al., 2015)]. Hence, we also sought to investigate whether there is an association between emotion characteristics of the movie and facial thermal changes.

## Materials and methods

### Thermal imaging and physiological data acquisition

Twenty healthy human participants (11 females, aged 22-30 years, mean = 25.7 years) were recruited for the study. All participants signed a consent form and were informed of the method. All participants had normal or corrected-to-normal vision. Exclusion criteria included habitual smoking, the presence of a chronic illness (e.g., cardiovascular or thyroid conditions), psychological disorders (e.g. depression or anxiety) or any other illness requiring regular medication. The study was approved by the Research Ethics Board of QIMR Berghofer and performed in agreement with the Declaration of Helsinki. The participants had the choice to withdraw from the study at any time. Each participant was compensated with a $50 voucher for their time. At the end of the data acquisition session a questionnaire was completed by the participants regarding subjective ratings of emotional response to the movie (Table S1). Physiological data from three out of twenty participants were excluded from analysis due to the failure of accurate trigger information and incomplete data acquisition. Questionnaire ratings could not be recorded from one subject. Informed consent to publish identifying images (RGB and thermal) was obtained from a subset of subjects.

Prior to the data acquisition, participants acclimatized for about 10 minutes within the experimental room. The temperature and the humidity of the room were kept within a steady range (22 ± 2 °C; 55–65% relative humidity). Where necessary, participants’ hair was secured away from their forehead with an unobtrusive hat. Participants were also asked to avoid alcohol and caffeine for at least 2 hours prior to the experimental session in order to minimize the vasoactive effects that these substances have on the skin temperature. Testing was performed exclusively between 2-5 pm to avoid any potential confounding effects of the circadian rhythm. After the participants had assumed a comfortable posture in a fixed chair, GSR and ECG electrodes were attached to fingers and arms respectively. A thermal imaging camera and an RGB video camera were then manually focused on the face. The researcher checked the recording quality and left the experimental room but retained audio-visual contact via CCTV. The paradigm consisted of a ∼20-minute short movie The Butterfly Circus (Weigel, Williams, & Weigel, 2009). Ten seconds of countdown was prefixed to the movie. Stimuli were presented on a 24-inch computer screen 40 cm in front of the subject. The sound was kept to the same levels for all subjects and was presented via two loudspeakers placed beside the stimulus screen.

An in-house built integrated hardware and software experimentation platform LabNeuro was used to integrate the multi-modal data acquisition. Thermal imaging of the face was performed by a FLIR A615 camera with a 15mm lens, an uncooled Vanadium Oxide (VoX) detector producing images of 640×480 pixels in size. The FLIR A615 provides a temperature detection range from -20 to 2000°C with the NEDT (noise equivalent differential temperature) smaller than 0.05°C at 30°C. The thermal camera response was blackbody-calibrated to nullify noise-effects related to the sensor drift/shift dynamics and optical artifacts. The sampling rate for thermal imaging was set at 5 Hz. This was performed in order to generate sufficient frames to balance out any potential movement artifacts by the participant. The cameras were incorporated into LabNeuro using the LabVIEW Image Acquisition library.

GSR recordings were obtained by two Ag/AgCl electrodes (0.8-cm diameter) filled with a conductive paste and attached to the distal phalanges of the index and ring fingers of the participant’s left hand. Skin conductance was recorded using a standard constant voltage system of 0.5 V and recordings were continuously digitized by an A/D converter with a sampling rate of 2 kHz. The recording of GSR data was acquired using National Instruments (NI) CompactDAQ modular IO hardware and software for the system was written using NI LabVIEW and NI Biomedical Toolkit. For ECG data collection, electrodes were attached to the mid-upper left arm, left wrist, and mid-upper right arm. We asked participants to put their hands-on chair hand-rests to minimize motion artifacts. ECG data was recorded with a sampling rate of 2 KHz. GSR signals and ECG data were collected simultaneously with the thermal imaging data.

### Extraction of facial thermal responses

The pipeline for this workflow is illustrated in Fig 1. Thermal video motion correction was achieved by applying a dense optical flow algorithm (Weinzaepfel, Revaud, Harchaoui, & Schmid, 2013). Specifically, the nonlinear motion vector field between each frame pair was first estimated by this method. Discontinuities in the transformation, which could distort the boundary of the face, were effectively removed by iterating motion correction and a Gaussian smoothing step applied to the motion vector field until convergence (where the displacement of all pixels is less than 1).

Similar to its use in fMRI data, spatial ICA was applied to identify and remove artifacts and to unmix statistically independent (true) sources of facial thermal signal change. To reduce noise and the number of parameters to be estimated, pre-whitening and dimension reduction were employed before the application of ICA. sICA was applied using a publicly available software package FastICA (http://research.ics.aalto.fi/ica/fastica/code/dlcode.shtml). The pipeline of the implementation of ICA together with motion correction is shown in Figure 1.

Though the motion correction and the ICA are both automated processes, the selection of physiologically meaningful independent components still needs manual inspection. To identify independent components with the most anatomico-physiological meaning we restricted attention to spatially localized components corresponding to broad anatomical boundaries such as nose, cheeks and forehead. The time series corresponding to each component was then normalized to have values between 0-1.

### Comparison of nose component signal with region-of-interest based nose-tip signal

A circle of 9-pixel radius was used to extract thermal signals from the nose-tip of thermal video data motion-corrected using optical flow (Fig 4A left). The time series were then normalized to have values between 0-1 for each subject to mitigate against spurious inter-subject differences. The correlation of ROI based nose-tip signal to that obtained from nose IC was computed, and significance of this correlation assessed using a non-parametric permutation test. Specifically, we calculated the group mean of the Pearson correlation between each subject’s nose component signal and their ROI based nose-tip signal. To generate the null distribution, each permutation used the ROI nose-tip signal with a random circular shift, so as to preserve temporal autocorrelation, and the group mean of the correlation between these shifted data and the thermal signals was computed. The null distribution was generated from 5000 permutation realizations.

**Figure 3.**
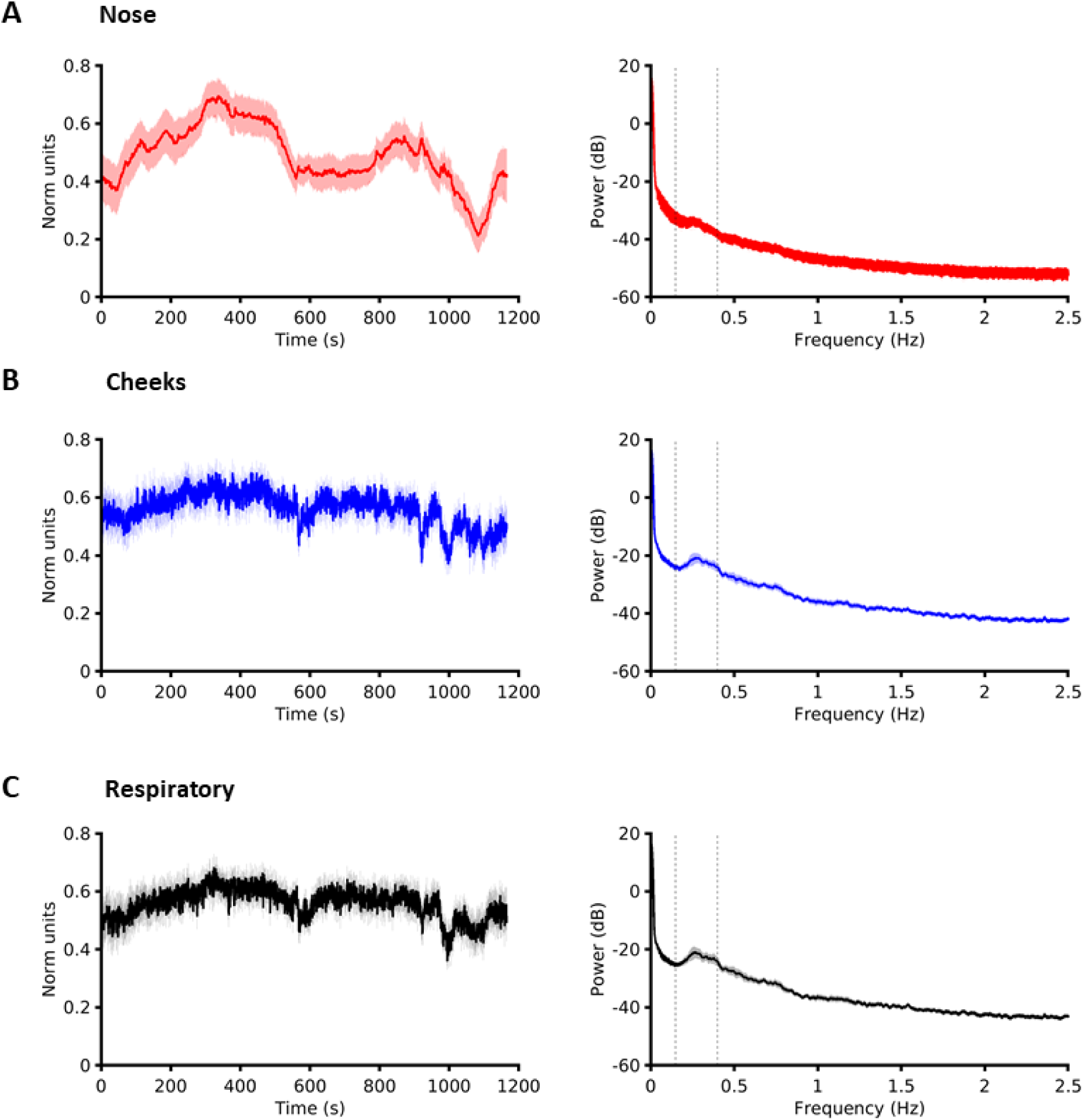
Group averaged component signals and their spectral signatures. **A)** nose component signal and its power spectra (right). **B)** bilateral cheek component signal and its power spectra (right). **C)** respiratory component signal and its power spectra (right). Vertical dashed lines on the spectral plots indicates the normal respiratory frequency range of 0.16 - 0.35 Hz. Respiratory and cheek components both seem to be affected by respiration whereas the nose component seems minimally affected by it. Shading indicates SEM.

**Figure 4.**
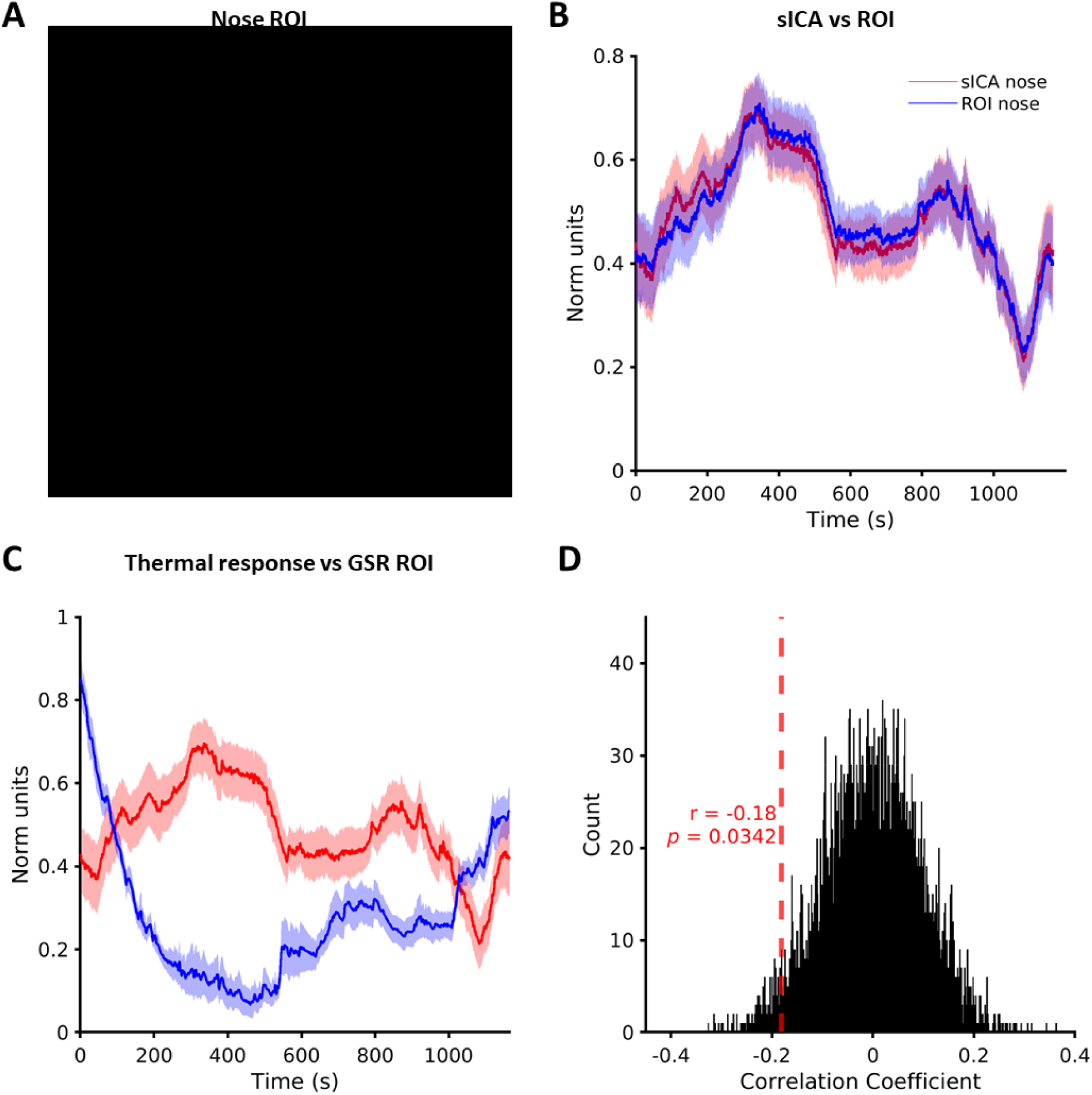
Nose component signal validation by comparison to ROI (region of interest) method and to GSR. **A)** ROI location on motion-corrected thermal image of a subject (L08) with nine-pixel radius. **B)** Comparison of group average nose sICA thermal component (red) with that of group average thermal signal obtained by ROI method (blue). **C)** Group average comparison of thermal response obtained from nose sICA component (red) to GSR (blue). Shading indicates SEM. **D)** Null distribution obtained with 5000 permutations showing statistical significance of negative correlation between thermal response and GSR. (Facial image in **A** not displayed as individual facial images display is against bioarxiv policies)

### Heart rate variability

The ECG signal was pre-processed using QRSTool software (Allen, Chambers, & Towers, 2007) to detect the R peaks with the ability to manually correct for missed peaks. Inter-beat interval (IBI) time series was then computed from this and normalised to have values between 0-1. R peak data were further analysed using HRVAS toolbox (Ramshur, 2010) to obtain heart rate variability (HRV) frequency domain measures. These were calculated via the auto-regressive method using a window size of 16 seconds, with 15 samples overlap, nfft of 1024 and cubic spline interpolation rate of 2 Hz (Sonkusare, Ahmedt-Aristizabal, et al., 2019). HRV data metrics were computed for the whole stimulus but edge effects for frequency estimation led to loss of approximately 8 seconds of data at the beginning and the end. Time-frequency decompositions of IBI are typically linked to autonomic influences in distinct frequency bands. Lower frequency (LF) HRV (0.04 – 0.14 Hz) mainly reflects changes in sympathetic and parasympathetic outflows, while high frequency (HF) variability (0.15 to 0.4 Hz) is primarily due to modulation of parasympathetic outflow (Akselrod et al., 1981; Pomeranz et al., 1985). Respiratory sinus arrhythmia (RSA) is a major contributor to HF HRV and is thought to be due to respiration modulating the cardiac vagal activity (Bernardi, Porta, Gabutti, Spicuzza, & Sleight, 2001).

### Comparison of nose component signal with skin conductance (GSR)

GSR is a standard measure of arousal. Event based studies have found an inverse relation between these two physiological measures (Shastri, Merla, Tsiamyrtzis, & Pavlidis, 2009; Sonkusare, Ahmedt-Aristizabal, et al., 2019). The GSR time courses were first detrended and low pass filtered at 5 Hz and subsequently normalized to have values between 0-1 for each subject to mitigate against spurious inter-subject differences in baseline. For statistically comparing GSR with nose component signal, identical permutation testing procedure as used for comparison of thermal signals to that ROI based signal was used.

### Inter-subject correlation of physiological signals

Pearson correlation coefficients between each pair of participants were computed separately for both thermal signals, GSR data and heart rate data. Inter-subject correlation (ISC) was computed as the mean pairwise correlation between participants. With 17 participants, each analysis comprised a total of 136 correlation pairs. Non-parametric permutation tests were used to identify statistically significant inter-subject correlation (p = 0.05) (Kauppi, Jääskeläinen, Sams, & Tohka, 2010; Nguyen et al., 2016). Specifically, the null distribution was generated from 5000 permutation realizations. These permutations comprised circularly shifted data in a way to preserve temporal autocorrelation in the physiological signals but disrupt correlations between subjects.

### Testing for similarity between two different recordings at group level

To assess statistically whether two different physiological responses were significantly correlated (such as the nose IC and nose-tip ROI thermal signals, the nose IC thermal signals with GSR, and the nose IC thermal signals with movie emotions), we employed non-parametric permutation testing. We first calculated for each subject the Pearson correlation coefficient between the two signals of interest. The correlation values were then averaged for all the subjects to obtain a group mean correlation value. To statistically test the significance of the obtained group mean correlation value, we applied non-parametric permutation tests with a null model where data from one modality was randomly circularly shifted relative to the other (thus preserving autocorrelation), before computing the group mean correlation of the null. Five thousand realizations were used to generate the null distribution.

***Figure 1.*** *The computational pipeline for employing spatial independent component analysis (sICA) on thermal imaging data* **(a)** Motion correction framework showing co-registered images with and without application of optical flow. For each subject, the whole thermal imaging data are aggregated into a single matrix, in which each row represents the thermal imaging data in one time point and each column stands for the time series of one single pixel **(b)** A mask to exclude background, i.e. neck and clothes, and retain only the face was applied to each frame. The data from these images were used as the source matrix **(c)** sICA - illustration of the mixing matrix, each column of which represents the time course of the corresponding source signal. An exemplar time series of nose components is shown. Each row of the source signal matrix represents one independent spatial map. Thermal image of subject L06 is used for illustrative purposes. (Facial images not displayed as individual facial images display is against bioarxiv policies)

### Facial expression analysis

Facial emotions expression scores, both for the movie actors and for the subject’s responses were obtained by processing the movie frames, and the RGB video recordings of the each subject through the Microsoft CNTK Face API (Cognitive Toolkit) (Del Sole, 2018). This software has been rated as one of the best for emotion categorisation (Khanal, Barroso, Lopes, Sampaio, & Filipe, 2018). The software takes a facial image as an input and returns the likelihood of each emotion class, across a pre-specified set of emotions, for each face in the image. The emotion classes permissible are anger, contempt, disgust, fear, happiness, neutral, sadness and surprise. They are ranked in descending order, with the overall total score summing to 1. The score was averaged for all the frames corresponding to each second. Briefly, emotion recognition is usually based on three key steps: 1) face detection, 2) feature detection—such as eyes and eye corners, brows, mouth corners, the nose-tip etc. and 3) feature classification—translation of the features into Action Unit codes, emotional states, and other affective metrics (Cowie et al., 2001). We summed the emotion scores for each emotion category for each subject before undertaking inter-subject correlation analysis to investigate the consistency of facial expressions among subjects.

## Results

Subjective ratings showed that the participants perceived the movie to be positively valenced (Table S1 Question 1: mean = 5.95, SD = 1.43) as well as emotionally intense (Table S1 Question 2: mean = 5.53, SD = 1.26).

### Consistent facial components identified by sICA

We extracted fifty independent components (ICs) from each subject. Of these, three components could be consistently identified across all subjects based on spatial and temporal characteristics. These were a nose component which mainly included pixels on the nose-tip, a cheek component which mainly contained pixels distributed on bilateral cheeks, and a respiratory component. The spatial patterns of these components clearly follow anatomical boundaries of facial features, supporting that they are driven by underlying physiological processes (Fig 2 left).

The temporal courses of these three ICs showed gradual evolution over the course of the movie (Fig 3 left). To investigate whether the nose and the cheek ICs identified were distinct from the respiratory component, we compared the power spectra of these signals (Fig 3 right). A frequency range of 0.16 - 0.35 Hz is indicative of normal respiration frequency (Russo, Santarelli, & O’Rourke, 2017). The nose IC was minimally affected by respiration whereas the power spectra of both the cheek and the respiration IC exhibited a peak at frequency range of 0.16 - 0.35 Hz. The similar time courses and power spectra of the bilateral cheeks to that of the respiration IC is consistent with the cheek IC’s diffuse spatial spread which likely includes respiration affected signals.

***Figure 2.*** *Distinct spatial components.* Representative components from one subject (L08) are shown. Three components were consistently identified in all subjects (except respiratory component absent in one subject). Color scale normalized between 0 and 1 for each component for display. Results from subject L09 are shown here. (Facial images not displayed as individual facial images display is against bioarxiv policies)

### Validation of sICA nose component by ROI tracking method

To validate the ICA-based method, we compared the time courses of the nose IC with those of thermal signals obtained using a ROI located at the nose tip (Fig 4A). Note that for thermal signal extraction with ROI method, we used the same motion correction algorithm based on optical flow. To our knowledge no other facial thermal imaging study has used an optical flow-based motion correction algorithm. The similarity between signals from the two methods was assessed by evaluating the group average Pearson correlation coefficient of nose IC and the thermal signal extracted from the corresponding ROI of the subject and statistically tested with non-parametric permutation testing (Fig 4B). (see Methods) (r = .94, p< 10-7, SI Fig S2 left).

### Inverse relation between nose component signals and GSR

GSR has been extensively validated as an arousal measure in psychophysiological studies (Kreibig, 2010). GSR thus provides a valuable benchmark to compare other psychophysiological measures and for facial thermal imaging this has previously been done, albeit for short event-based stimuli (Pavlidis et al., 2001; Shastri et al., 2009). We thus compared the time courses of the nose component with that of corresponding GSR signals to further investigate their response relationship during naturalistic emotional experiences. Permutation testing showed a significant inverse relationship (r = -.18, p = .03) (Fig 4C, D).

### Similarity of physiological changes across subjects

We then examined the consistency of all the corresponding physiological variables across subjects using inter-subject correlation (ISC) analysis. We found significant consistency between subjects in the dynamic thermal response patterns for the nose IC (mean r = 0.12, p = .007, Fig 5A), and still greater consistency for dynamic GSR (mean r = 0.50, p < .0001, Fig 5B). The high ISC of GSR signals seemed to be mostly attributable to the gradual decrease of GSR at the beginning of the movie viewing – by contrast the ISC of GSR signals during the second half of the movie was 0.08 (p = .03), far lower than the ISC coefficient value for the thermal signal in the first half of the movie of .14 (p=.003). ISC of frequency components of heart rate variability (HRV) also showed significant inter-subject correlation (low frequency HRV: mean r = 0.03, p = .0002, Fig 5C left, high frequency HRV: mean r = 0.03, p = .00004, Fig 5C right, pFDR =.05).

**Figure 5.**
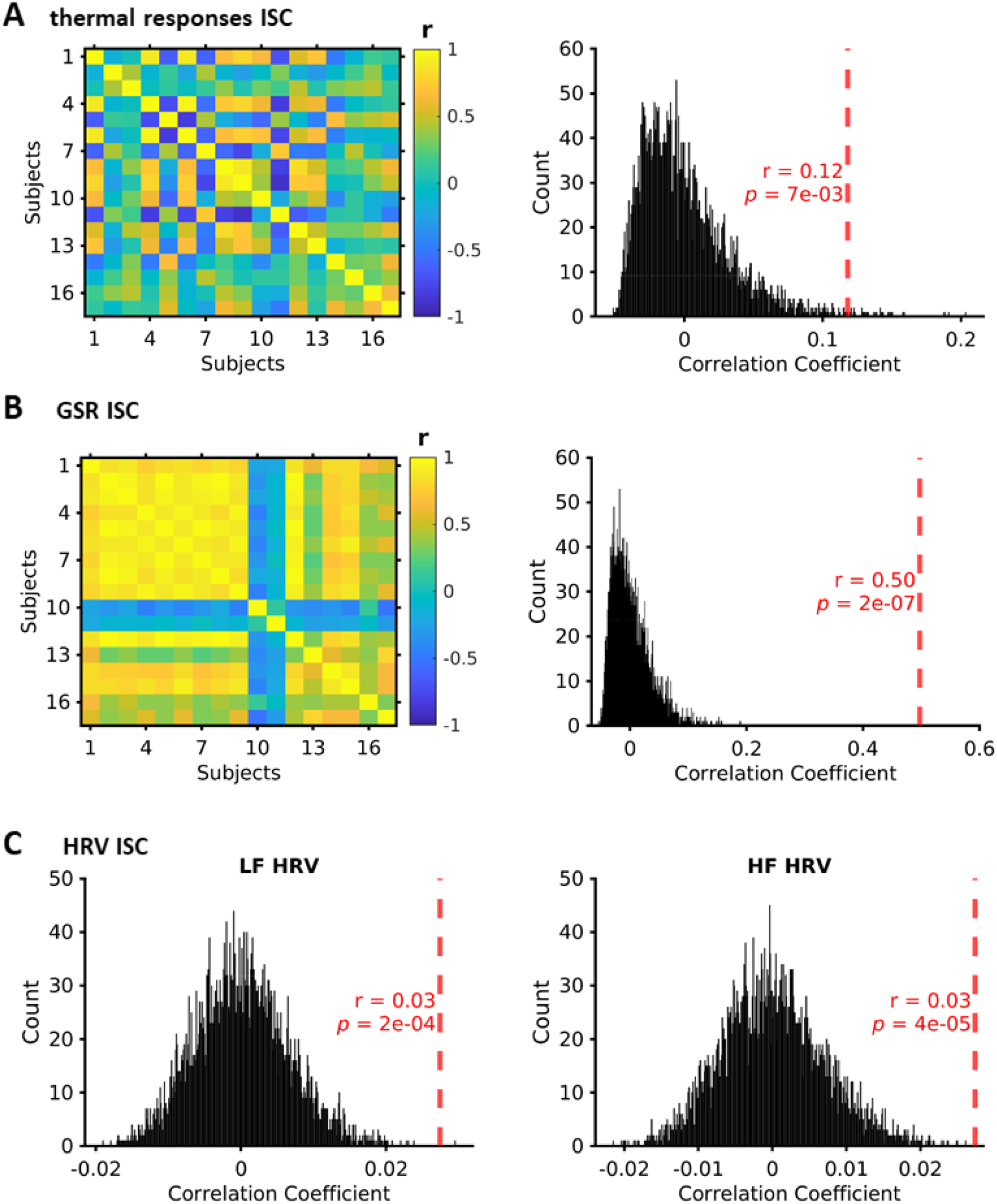
Inter-subject correlation (ISC) analysis. **A)** correlation matrix of nose component thermal response time series (left) and null distribution obtained with 5000 permutations (right) (see methods) showing statistical significance of positive mean correlation shown in red. **B)** correlation matrix of GSR responses curves (left) and right - null distribution obtained with 5000 permutations (right) (see methods) showing statistical significance of positive mean correlation shown in red **C)** ISC analysis of low frequency heart rate variability (LF HRV) and its statistical significance (shown in red) (left) across participants when tested with 5000 permutations. ISC analysis for high frequency (HF) HRV (right). FDR corrected for statistical comparisons for LF and HF HRV. Colour bar r denotes Pearson’s correlation coefficient. Histogram r denotes mean correlation coefficient at group level.

### Thermal correlates of emotion during movie viewing

We then evaluated whether thermal responses were correlated with the emotional content of the movie. Previous studies, using an event-related design, have found the latency of thermal responses to be around 3.8 s (Merla & Romani, 2007) and 4-5 s (Sonkusare, Ahmedt-Aristizabal, et al., 2019) after stimulus onset. In this study we employed a naturalistic paradigm with continuous narrative instead of discrete events, hence, to compare the emotions of the movie and thermal signals we correlated a 3 second lagged thermal signal with the movie emotion time series. The emotional content of the movie was quantified by computing the scores of facial emotional expressions of the actors in each frame of the movie averaged for each second (see Methods). Thus, each second of the movie was assigned an emotional score according to the emotion detected for that frame (happiness, sadness, surprise, anger, fear, disgust and contempt), excluding the neutral category (Fig 6A). A non-parametric permutation test revealed a significant negative correlation between the thermal signal and happiness scores (r = 0.06, pFDR = .0004, Fig 6 B) and a significant positive correlation between thermal signal and anger scores (r = 0.05, pFDR = .004, Fig 6 B). These results are also replicable when using a 4 second lagged thermal signals (happy r = .06, pFDR = .0001, anger r = .05, pFDR = .004).

**Figure 6.**
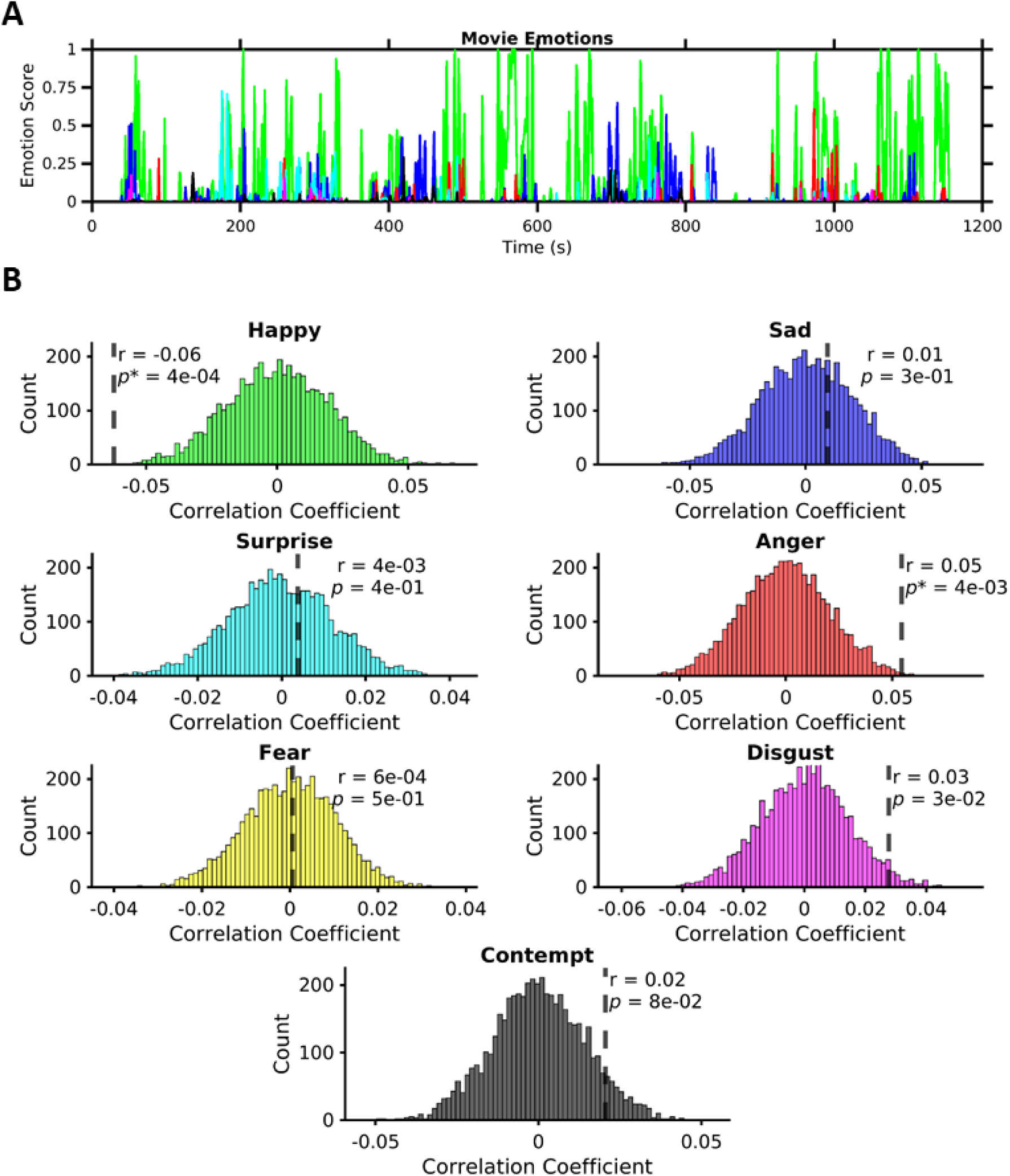
Thermal correlates of emotion. **A)** The averaged emotion scores from facial expressions detected in movie frames. **B)** Plots with coloured histograms represent the corresponding emotion category. Histograms were obtained from permutation testing to test statistical significance of correlation between different emotion scores and nose thermal signals. Thermal signal correlation with happy and anger emotion are statistically significant after multiple comparisons (*p*\* denotes significant results with p_FDR_ < .05). Scientific notation used for r and *p* values where appropriate.

### Subjects’ facial expression of emotion was minimal

We also recorded facial video data of subjects watching the movie. The subjects minimally facially expressed the emotions with mean emotional scores for happy and sad categories predominantly below .1 score (SI Fig S1). There was no significant inter-subject correlation for facial expressions (all emotions summed for each subject) across subjects (r = 0.02, p = .08, SI Fig S2 right).

## Discussion

We sought to develop a new data-driven method for facial thermal imaging analysis and determine whether long term facial thermal fluctuations elicited by a dynamic, naturalistic stimulus convey emotional signatures. We validated our method with traditional ROI analysis showing the robustness of our approach. We also uncovered the association of changes in the nose-tip thermal signal with dynamic emotional changes in the stimulus. We further demonstrated common physiological responses across subjects as measured by thermal signals, heart rate metrics and GSR, underlining common changes in emotional states induced by the naturalistic stimulus. Crucially these results were obtained in an ecologically valid context.

We validated our methodology by comparison to the traditionally employed ROI method and demonstrated remarkably similar results for nose responses with both these methods. The robust correspondence of results with the two methods can be due to state-of-the-art motion correction technique employed. As far as we are aware no other thermal imaging study has utilized optical flow for motion correction of thermal video. Existing thermal imaging studies employing motion correction have used spatial cross-correlation (Manini et al., 2013; Pinti et al., 2015) but these are of limited effectiveness when used with a 3D surface that is not flat, such as the human face. The successful applications of optical flow and spatial ICA in this work thus provide two separate novel methodological contributions, as the former can be used to provide motion correction for ROI based methods as well.

The spatial components of nose and cheeks identified in this study correspond to facial regions widely studied in previous studies (Ebisch et al., 2012; Merla & Romani, 2007; Nakanishi & Imai-Matsumura, 2008; Pavlidis et al., 2001; Shastri et al., 2009). In addition to these, we consistently identified a respiration related component. Respiration monitoring via thermal imaging has already been demonstrated (Cho, Bianchi-Berthouze, & Julier, 2017; Cho, Julier, Marquardt, & Bianchi-Berthouze, 2017). The different facial components were associated with distinct thermal responses. Specifically, the cheek component and respiratory component showed similar time courses and with both being affected by respiration noise. Moreover, the spatial maps of the cheek component showed a diffuse distribution and hence was likely noisier. The successful application of video motion correction and spatial ICA in this work demonstrates the utility of this approach for extracting robust thermal signals in longer-time naturalistic paradigms. A promising avenue of future research will be to use spatio-temporal ICA in place of spatial ICA (Stone, Porrill, Porter, & Wilkinson, 2002) to attempt to separate sympathetic and parasympathetic influences in the same facial regions.

Facial blood flow changes detected by thermal imaging are caused not only by sympathetic (predominantly) but also parasympathetic influences (Drummond, 1994; Segade & Sua, 1990). Thus, it is a more complex signal in comparison to GSR which is a uni-dimensional sympathetic response representing arousal (Bach, Friston, & Dolan, 2010; Boucsein & Hoffmann, 1979). Thermal signals thus seem to convey more nuanced representation of sympathetic-parasympathetic balance. The more complex nature of thermal signals may also contribute to greater subject to subject variability and may explain the unexpected anti-correlations between a few subjects (Fig 6B). GSR signals are considered the gold standard in peripheral neurophysiological and psycho-physiological studies, providing a benchmark for validation in IRI research. Overall, these results further highlight the biological underpinnings that might make thermal responses useful in differentiating emotional valence.

All the physiological variables (namely thermal imaging, heart rate metrics and GSR) showed significant similarity among subjects, highlighting the ability of a dynamic emotional stimulus to induce common emotional states. This can be attributed to the use of naturalistic stimuli, which have been argued to better evoke physiological responses than traditional stimuli (Gross & Levenson, 1995). Naturalistic stimuli offer a trade-off between completely uncontrolled stimuli and unconstrained conditions on the one hand (for instance resting state paradigms in neuroimaging), and the strict control of simplistic stimuli (for instance auditory odd-ball paradigm), placing relevant ecological constraints on physiological processes (Sonkusare, Breakspear, et al., 2019). As emotion is built over a longer narrative, these stimuli can simulate common, everyday emotional experiences and evoke robust and consistent responses in subjects as evident from our inter-subject correlation analysis. These analytical methods of inter-subject correlation analysis, when applied to neuroimaging, have also shown shared neuronal processes underlying emotional states (Hasson et al., 2004; Nguyen et al., 2016; Pajula et al., 2012). Thus, the consistent responses across subjects reported in our study for diverse physiological variables provide convergence of findings on a body and brain level and reinforce the wider use of naturalistic paradigms to comprehensively investigate human emotions. In similar vein though we did not find a temporal consistency of facial expressions among subjects (Fig S2). Facial emotion displays encountered in everyday life situations show high variability (Nummenmaa & Calvo, 2015; Scherer & Ellgring, 2007) and spontaneous expressive behaviour is more complex and ambiguous (Pantic, 2009). This was true in our subjects as well, as they had low emotion expression scores.

The movie stimulus we employed is an engaging film with various emotional undertones (Nguyen et al., 2016). Although one-to-one mapping of facial emotion content in the movie and emotion induced in the viewers is complex, the former can act as a proxy measure to quantify emotional undertone of the movie. Importantly, the correlated nature of the two for our stimulus is signified by the majority of the movie frames (Fig 4 top panel) with a high happy emotion score. We demonstrated that facial thermal response is negatively correlated to this happy state. Anticorrelation of thermal responses with positive emotions has been previously demonstrated in a controlled paradigm (Matsukawa et al., 2017). However contrasting results have also been found (Zenju et al., 2004) but which have used static images as stimuli. Our findings were obtained in a naturalistic context. Furthermore, another emotion, anger, was positively correlated with thermal responses. Anger and its association with increased facial colour change is well documented (Changizi, Zhang, & Shimojo, 2006). Many studies also support the association of the colour red with anger (red facilitating the identification of anger expressions) (Drummond & Quah, 2001; Young, Elliot, Feltman, & Ambady, 2013). Even the English language is replete with phrases such as “red with anger” depicting increased blood flow to the face and hence increased temperature. Aside from these significant findings, we did not find any association with other emotion categories. Further research to parsimoniously characterize the relationship between emotion differentiation and temperature variations of the face is thus needed. Still, a single autonomic dimension may not be sufficient to comprehensively characterize all different emotions [(Kreibig, 2010) however also see (Rainville et al., 2006)]. Hence, newer techniques such as thermal imaging providing contact-less recordings in ecological valid contexts may compliment traditional measures and others techniques less commonly used (such as contact-based measures: systolic and diastolic blood pressure, respiratory measures, finger temperature) to capture various aspects of the autonomic system or improve the accuracy of existing ones in differentiating emotions.

For the thermal responses and facial colour changes generated in the experience of emotion, there is evidence that these supplement communication of emotions (Alkawaz, Mohamad, Saba, Basori, & Rehman, 2015). Just as thermal imaging captures these facial blood flow changes, some of this information is also available to human vision. Indeed, emotion dependent facial blood flow changes were visually interpreted by observers and have also been found to convey information partially independent from that conveyed by facial expressions (Benitez-Quiroz, Srinivasan, & Martinez, 2018). In some facial thermal imaging studies, facial movements and expression may confound the thermal signal. However in this study the emotional information conveyed in thermal responses is unlikely due to facial movements based on three main reasons: 1) we used a nose-tip component signal which is minimally affected in facial expressions, 2) the subjects minimally expressed facial emotions and 3) we removed facial movement caused by changes of facial expression by using a robust motion correction algorithm. Thermal responses thus can be assumed to capture physiological phenomenon aiding in emotional experience and communication.

## Limitations of the Study

Some caveats and limitations of this study should be noted. First, the three consistent ICA components were identified manually based on visual inspection. An objective automated component selection method can help overcome this limitation in the future (Salimi-Khorshidi et al., 2014). Motion correction of facial video data has inherent challenges due to the 2D nature of the data while the actual face is a 3D object. We obtained motion correction using 2D dense optical flow, while future work may explicitly map 2D thermal data to a 3D facial model to achieve more precise motion correction. A common standard space for registration of all subjects’ faces will also improve the spatial decomposition. Another limitation is the lack of a control condition or resting state data to unequivocally conclude that the physiological changes were emotionally driven. While there was significant correlation to changes in emotion content of the movie, other cognitive factors may also have contributed to the physiological changes. Finally, for the correlation between emotion and thermal response, we note that while significant, the correlation coefficient values are not high. This is not unexpected as the automated scoring of emotion content was based on discrete facial emotion categories, which may be limited in capturing the complex nature of emotions portrayed in a movie. Future studies could compliment naturalistic stimulus with separately acquired continuous subjective ratings of valence and arousal to further probe temperature perturbations dependence on emotions.

In sum, we developed a novel data-driven analytical technique for facial thermal imaging analysis and showed robust facial thermal changes evoked by an emotional movie which were correlated with the emotional content of facial expressions in the movie. We also uncovered a physiological consistency among subjects thus signifying common responses elicited by the movie. Future studies could use simultaneous thermal and brain recordings such as combined facial thermal imaging with functional near infrared spectroscopy (Pinti et al., 2015) or electroencephalography (EEG), to investigate the brain-body interaction.

## Data and Code Availability

Code related to this paper is available from the authors. Data used in the study includes personal identifying facial images of participants, and local ethics approval mandated strict privacy restrictions around their availability outside of the named investigator team. Researchers wishing to access these data will require local ethics approval and a data sharing agreement with QIMR Berghofer.

## Acknowledgments

None

## Author Contributions

Conceptualization, C.G., S.S., V.N.; Data Curation, S.S., V.N.; Methodology, M.J.A., T.P., S.S.; Investigation, S.S., M.J.A., T.P.; Writing – Original Draft, S.S., T.P., Writing – Review & Editing, S.S., V.N., T.P., M.B., S.F., C.G., M.J.A.; Funding Acquisition, M.B., C.G.; Resources, S.F., M.B.; Supervision, M.J.A., C.G., M.B.

## Declaration of Interests

The authors declare they have no competing interests.

## Supplementary Material

**Table S1.**

|  | Questions probing subjective emotion ratings in the questionnaire | Ratings |
| --- | --- | --- |
| 1 | Is your emotional reaction to the film positive or negative overall?<br>(**1** = *extremely negative*, **4** = *neutral*, **7**= *extremely positive*) | **Mean = 5.95, SD = 1.43** |
| 2 | Overall, was your emotional reaction to the film calm or intense?<br>(**1** = *extremely bored/calm*, **4** = *neutral*, **7** = *extremely excited/intense*) | **Mean = 5.53, SD = 1.26** |
| 3 | Which emotion best describes what the film made you feel?<br>'happy', 'surprise', 'sad', 'disgust', 'fear', 'anger', 'others' | **14-happy, 4-surprise, 1-sad** |
| 4 | How strongly did you feel this emotion during the film?<br>(**1** = *Not at all*, **4** = *mildly* **7** = *extremely*) | **Mean = 5.79, SD = 1.13** |

SD – standard deviation

**Figure S1.**
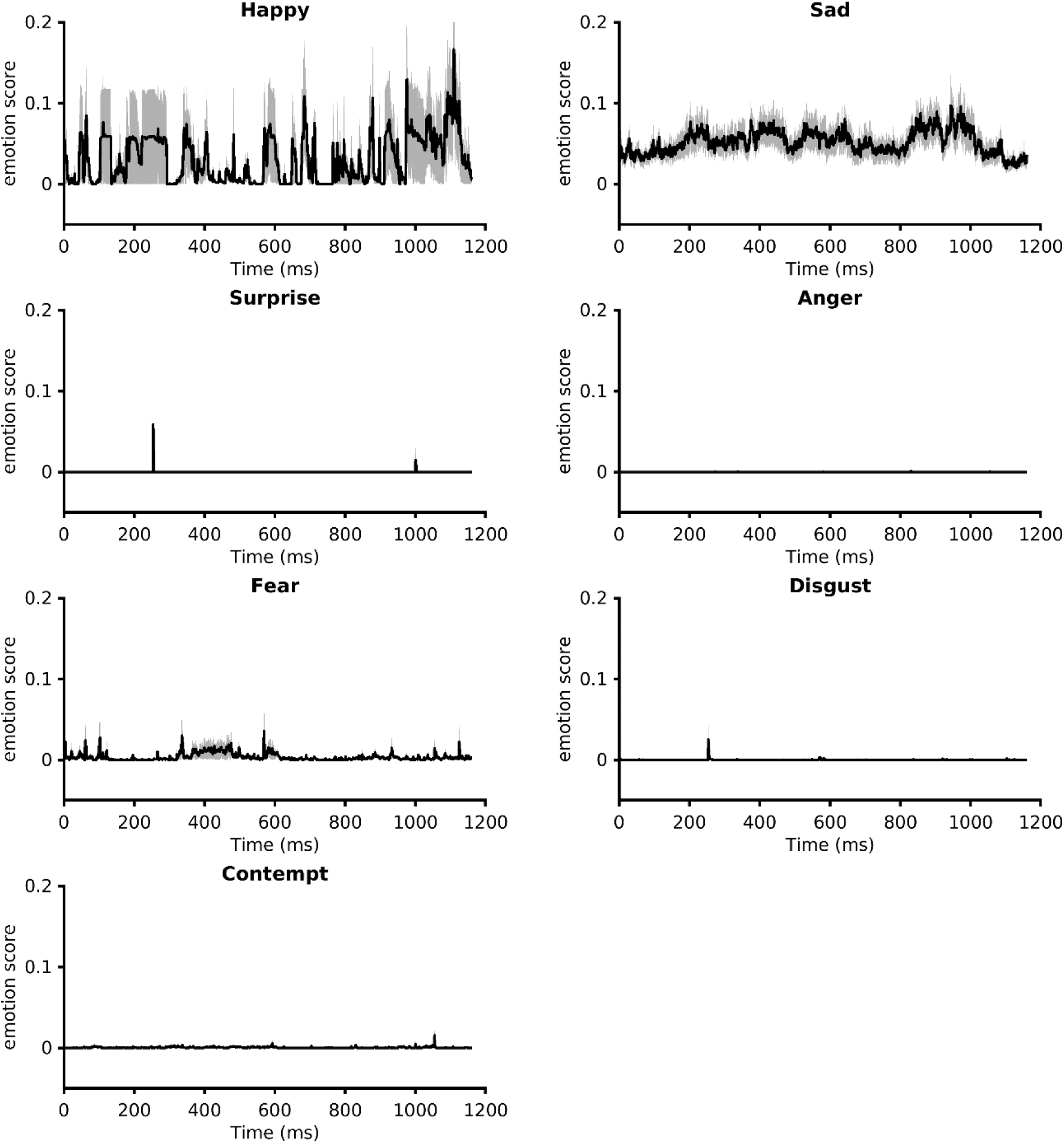
Facial expression scores of participants for various emotions. These plots show the mean facial expression scores with shading denoting sem. The emotion scores were predominantly below a score of .1.

**Figure S2.**
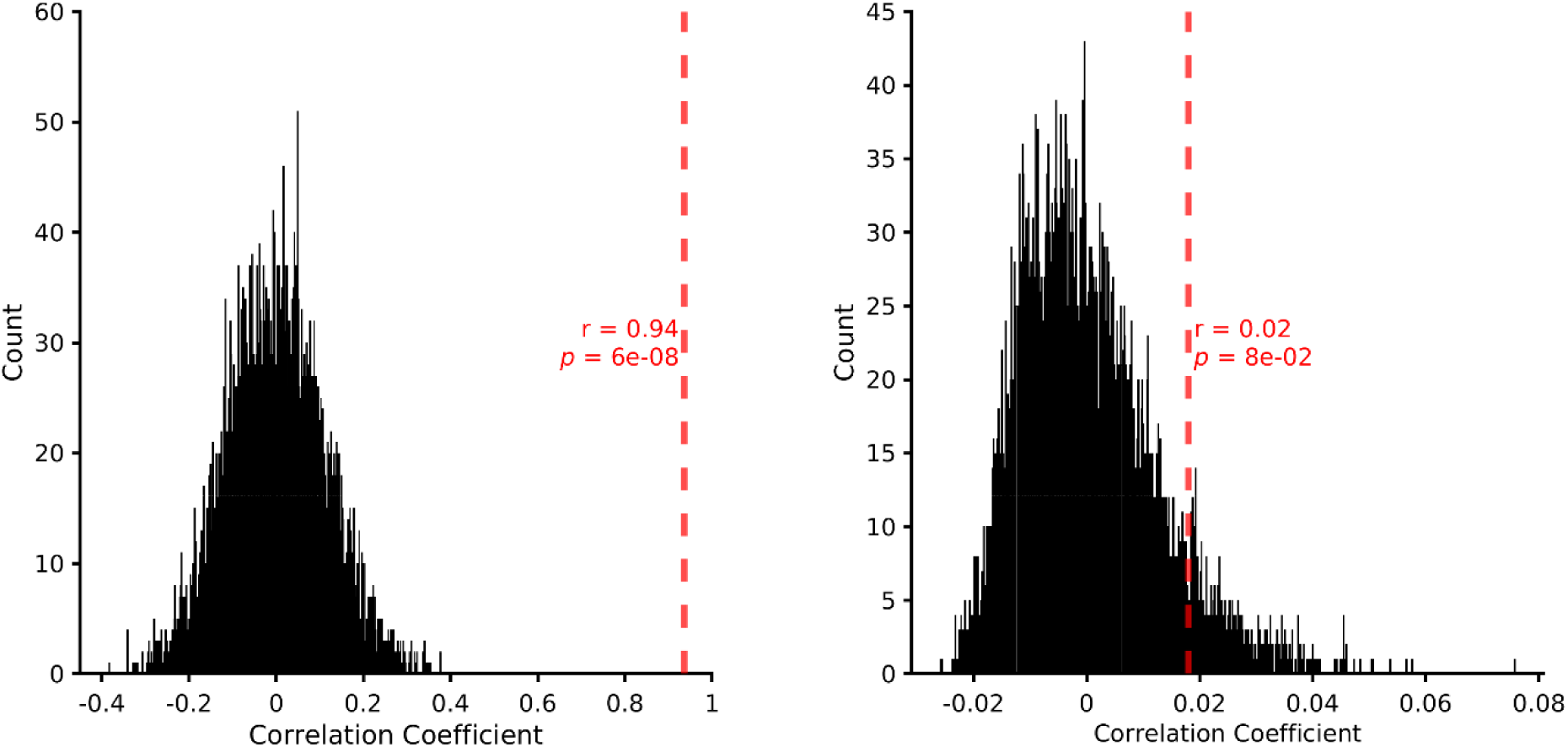
Correlation of sICA nose component with ROI nose signal, and Inter-subject correlation analysis for facial expressions. *Left*: null distribution obtained with 5000 permutations (see methods) showing statistical significance of positive correlation between nose thermal response from sICA and ROI method, *right*: null distribution obtained with 5000 permutations (see methods) showing no statistical significance of inter-subject correlation for subjects’ facial expressions.

***Figure S3.*** *Spatial components when 80 components were retrieved, instead of 50*. Representative components from one subject (L08) are shown. Note cheek component decomposes into two separate components. Color scale normalized between 0 and 1 for display. (Facial images not displayed as individual facial images display is against bioarxiv policies)

## Notes

### Competing Interest Statement

The authors have declared no competing interest.

